# Optic nerve intraneural stimulation allows selective visual cortex activation

**DOI:** 10.1101/311035

**Authors:** Vivien Gaillet, Annarita Cutrone, Paola Vagni, Fiorenzo Artoni, Sandra Alejandra Romero Pinto, Dario Lipucci Di Paola, Silvestro Micera, Diego Ghezzi

**Affiliations:** Medtronic Chair in Neuroengineering, Center for Neuroprosthetics and Institute of Bioengineering, School of Engineering, École Polytechnique Fédérale de Lausanne, Switzerland; The BioRobotics Institute, Scuola Superiore San’Anna, Italy; Bertarelli Foundation Chair in Translational Neuroengineering, Center for Neuroprosthetics and Institute of Bioengineering, School of Engineering, École Polytechnique Fédérale de Lausanne, Switzerland

## Abstract

Retinal prostheses have been developed to restore a functional form of vision in patients affected by outer retinal layer dystrophies. Other approaches, namely optic nerve, thalamic, and cortical prostheses are under investigation to expand this toolbox both as clinical devices and as tools for fundamental research. Optic nerve stimulation is attractive since it bypasses the entire retinal network and directly activates nerve fibers. At the same time, it still takes advantage of the high-level information processing occurring downstream in the visual cortex. Here we show that a new intraneural electrode array (OpticSELINE) is effective in inducing the activation of the visual cortex upon electrical stimulation of the optic nerve. We also demonstrate that intraneural optic nerve stimulation induces selective cortical activation patterns depending on the stimulating electrode, thus suggesting that the OpticSELINE possesses spatial selectivity in fiber stimulation. In conclusion, the OpticSELINE can be used both as visual prosthesis for blind patients and as tool to further investigate the effect of the electrical stimulation in the visual system.

---

Visual prostheses recently emerged as tool to fight blindness; a medical condition affecting more than 30 million people worldwide^1^. Starting from early pioneering works^2,3^, over the past 50 years several types of visual prostheses have been proposed and classified by their location along the visual pathway^4,5^, including: subretinal^6^, epiretinal^7^, suprachoroidal^8^, optic nerve^9^, thalamic^10^, and cortical prostheses^11^. Retinal implants quickly became the preferred strategies, since they can benefit from the natural information processing along the visual pathway, despite the limit to treat only diseases affecting retinal photoreceptors^12^. In various clinical trials, retinal prostheses demonstrated the capability to restore a functional form of vision^13,14^, and today several research groups are developing novel retinal prostheses^15–17^. On the other hand, cortical prostheses are still facing several technical and biological challenges; the invasiveness of surgical implantation, the risk of focal seizures induced by direct cortical stimulation, and the requirement of high-level information coding pose serious concerns for their clinical application^12,18^. On the contrary, optic nerve stimulation is attractive since it bypasses the entire retinal network and directly activates nerve fibers. At the same time, it still takes advantage of the high-level information processing occurring in the visual cortex. Optic nerve stimulation has been pioneered with the implantation of a 4-contact epineural electrode in the intra-cranial trait of a blind subject^19^, in which electrical stimuli have been able to elicit localized phosphenes. After few months of training and psychophysical testing, the patient has been able to distinguish line orientations as well as shapes and symbols despite using only 4 electrodes. Another patient has been implanted later with a 8-contact electrode^20^, confirming the possibility of restoring functional vision by using optic nerve stimulation^21–23^. However, in those trials the induced phosphenes have been reported as irregulars; the use of epineural electrodes may be the cause^24^, due to their limited mechanical stability. Following this approach, an optic nerve prosthesis is currently under investigation by the C-Sight project, which is testing the stimulation of the intra-orbital region with a 4-filament electrode; acute results have been documented in rabbits^25,26^ and cats^27^. This project employs penetrating platinum-iridium electrodes, which are characterized by high stiffness inducing a large mechanical mismatch with the nerve; in turn this may have consequences for chronic implantation. On the contrary, transverse intra-fascicular multichannel electrode (TIME) arrays, micro-fabricated via thin-film technology, have already demonstrated their superior capability in several applications related to nerve stimulation^28,29^. Indeed, intraneural electrodes show a higher selectivity in fiber stimulation^30,31^ with respect to epineural electrodes. More recently, a self-opening intraneural electrode (SELINE) demonstrated its improved mechanical stability^32^ and biocompatibility over a period of 6 months^33^. In this work, a modified version of the previously described SELINE electrode array^32,33^, has been successfully exploited as visual prosthesis based on optic nerve stimulation.

## Results

### Electrode design and characterization

The OpticSELINE is a polyimide-based looped structure with a total length of 33 mm, a maximum width of 3 mm, and an overall thickness of 0.012 mm (Fig. 1a). A polyimide-based extension flat cable of 35 mm allows the connection between the electrode and the head-plug connector. It has twelve stimulating electrodes (six electrodes per side, area of 0.008 mm^2^) plus a reference and a ground electrode outside the active area. Each side has two three-dimensional flaps that extend from the main body and carry two electrodes each; two more electrodes are located on each side of the main body. The width of the active area is 0.43 mm and the length is 1.25 mm. Each flap has a width of 0.15 mm and a length of 0.48 mm (Fig. 1b). Four alignment bars (width of 0.1 mm) have been included to ease the insertion procedure and verify that the active area is located inside the optic nerve (Fig. 1c). The OpticSELINE has been designed in agreement with the anatomical structure of the rabbit’s optic nerve, which has an average (± s.e.m.) diameter of 1.45 ± 0.04 mm (**Supplementary Fig. 1**).

**Figure 1.**
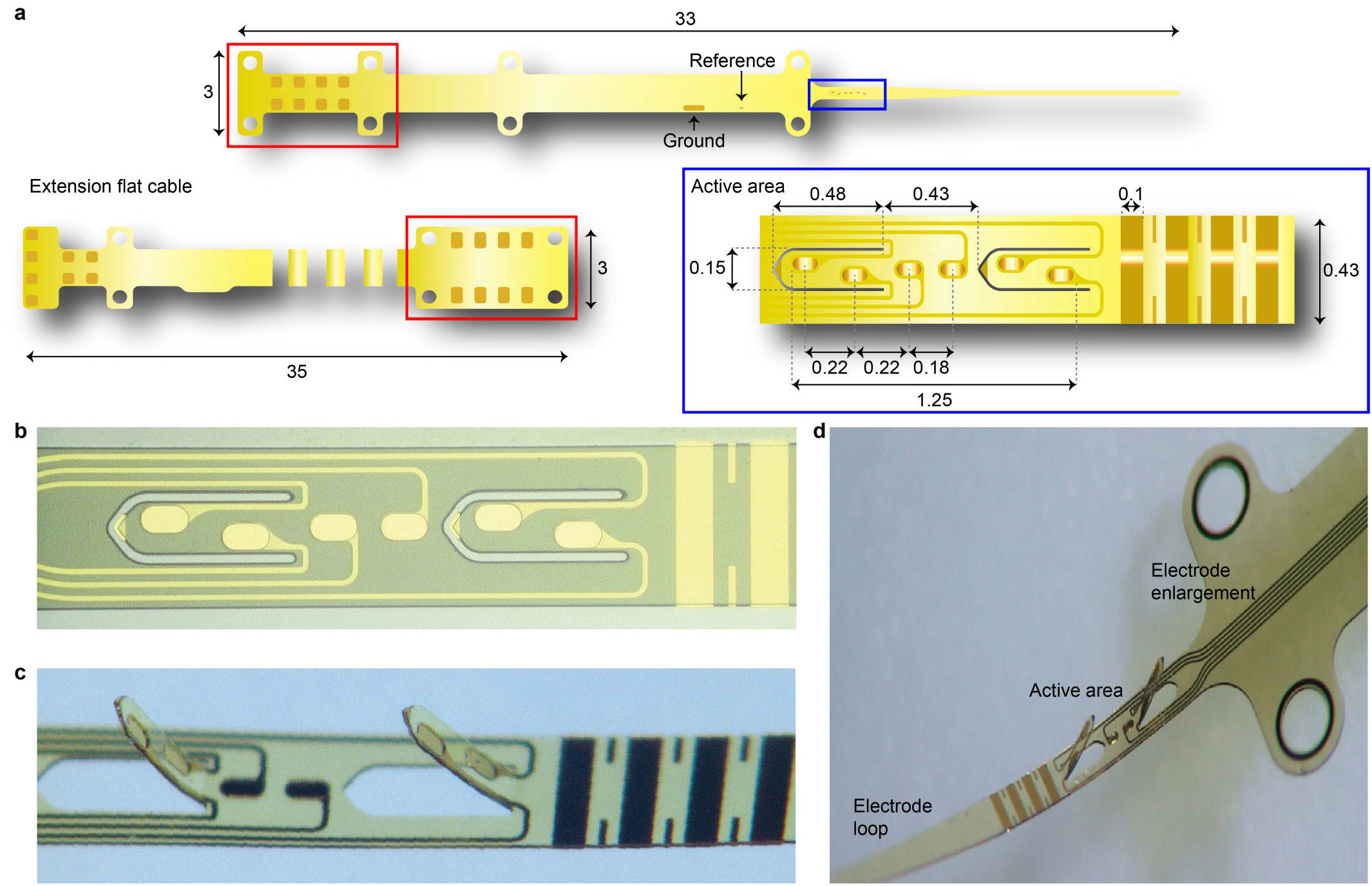
OpticSELINE design and fabrication. **a**, Sketch of the OpticSELINE. The red box highlights the connection area between the electrode and the flat cable terminated with an Omnetics connector (not shown). The blue box highlights the active area where 2 flaps and 6 electrodes are visible. Dimensions are in mm. **b**, Magnification of the active area after microfabrication. **c**, Enlarged view of the flaps, electrodes, and alignment bars after three-dimensional shaping. **d**, Picture of one side of the OpticSELINE. The electrode enlargement is used as a stopper to avoid excessive insertion of the array within the nerve.

The OpticSELINE has been characterized electrochemically and mechanically. First, cyclic voltammetry has been performed to determine the charge storage capacity of the electrodes (Fig. 2a). This test has been executed before and after an accelerated ageing time equivalent to an implantation time of 6.4 months (6 days at 87 °C)^34^. The mean charge storage capacity is not significantly different before and after ageing (Fig. 2b; *n* = 18; p = 0.1919, paired t-test). Given the small electrode area (0.008 mm^2^), this value of charge storage capacity is comparable to values found in literature for gold microelectrodes^35^. Similarly, the impedance spectroscopy has been performed (Fig. 2c), and the mean magnitude at 1kHz has shown non-significant statistical difference before and after ageing (Fig. 2d; *n* = 30; p = 0.7613, paired t-test).

**Figure 2.**
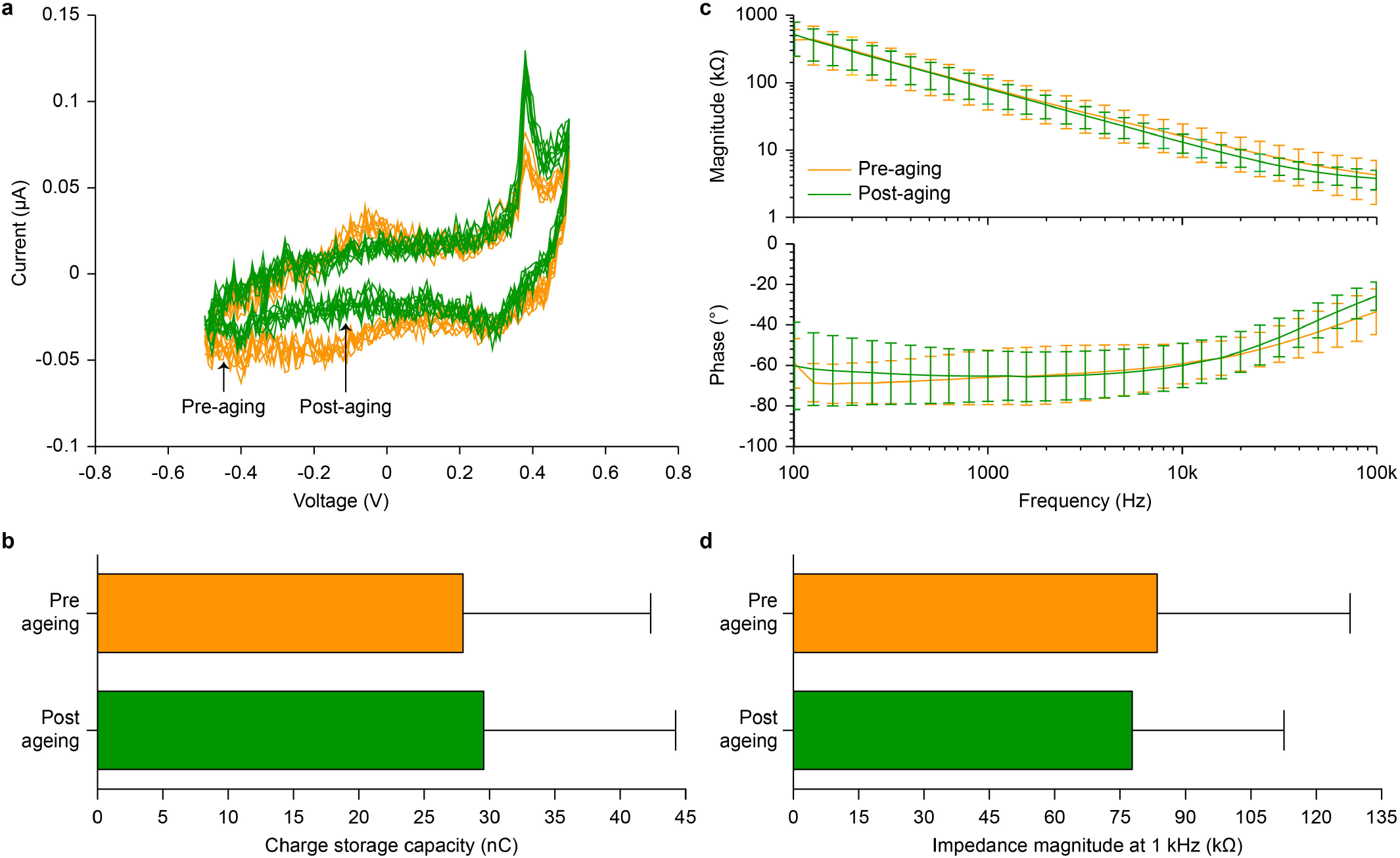
Electrochemical characterization. **a**, Cyclic voltammetry performed on the electrodes before (orange) and after (green) accelerated ageing. Representative example of 1 electrode with an overlay of 10 repetitions. **b**, Quantification of the mean (± s.d.) charge storage capacity (*n* = 18). **c**, Mean (± s.d.) magnitude (top) and phase (bottom) of the electrode impedances before (orange) and after (green) accelerated ageing (*n* = 30). **d**, Quantification of the mean (± s.d.) impedance magnitude at 1 kHz.

A mechanical characterization has been performed to verify the compatibility of the electrode array with insertion forces and the stability within the nerve. During insertion experiments (Fig. 3a) two major force peaks have been observed (Fig. 3c,e) corresponding to the insertion of the loop inside the nerve (peak 1) and to the entry of the enlarged area of the device (peak 2). The stability of the electrode within the nerve has been evaluated with an extraction experiment (Fig. 3b). During extraction two major force peaks have been observed (Fig. 3d,f) corresponding to the force necessary to extract the three-dimensional flaps (peak 3 and 4) followed by a flat phase relative to the slippage of the loop through the nerve (5). Measured extraction forces are 25 folds larger than the forces required to extract an electrode array without three-dimensional flaps, as previously measured in the rat sciatic nerve^32^. This confirms that the presence of three-dimensional flaps enhances the anchorage of OpticSELINE within the nerve and provides a better mechanical stability.

**Figure 3.**
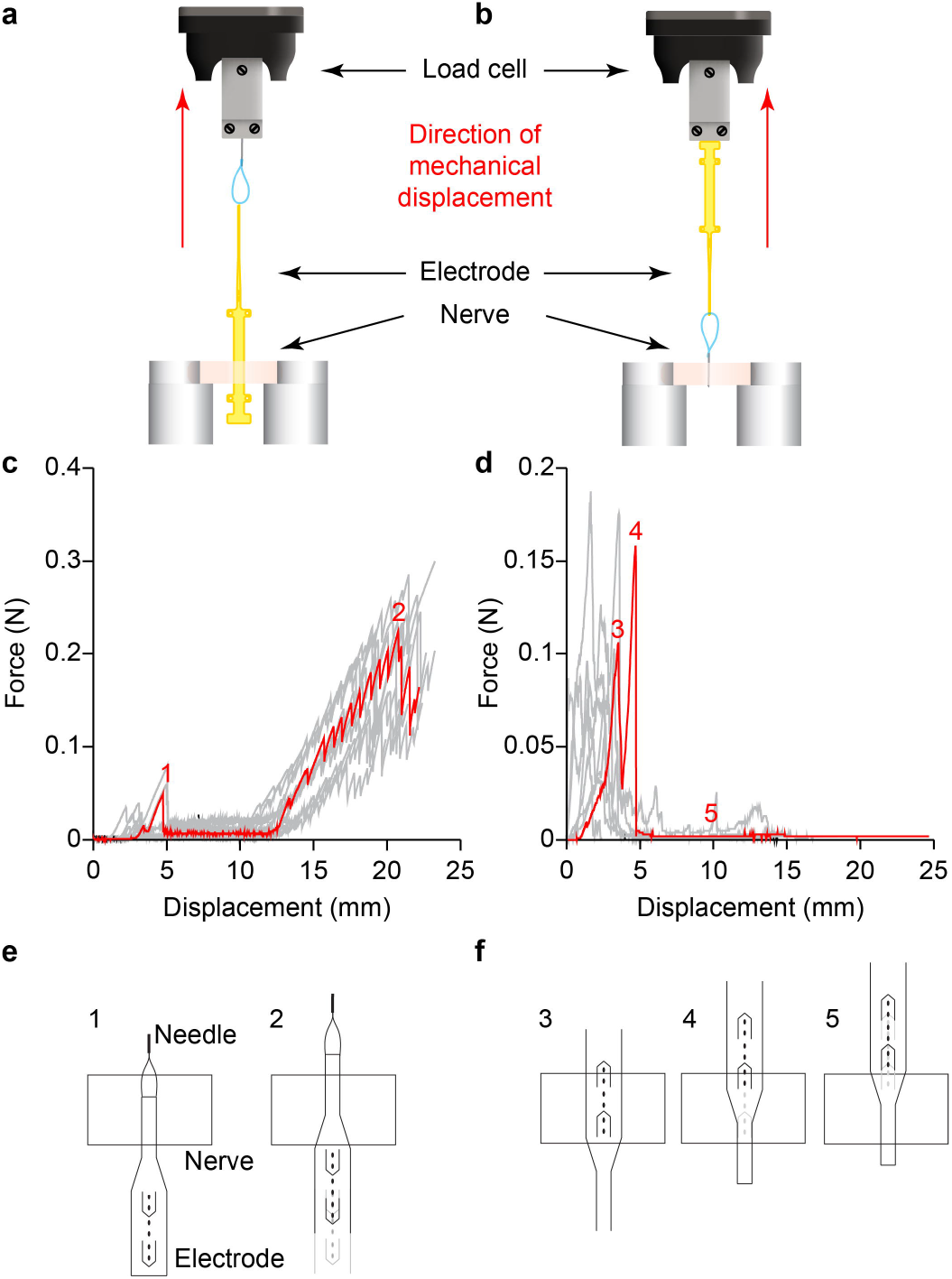
Mechanical characterization. **a,b**, Scheme for the ex-vivo insertion (**a**) and extraction (**b**) experiments in explanted optic nerves from New Zealand White rabbits. **c**, Forces during insertion in the optic nerve. Several insertion trials (*n* = 10) are shown in grey while, in red, a selected example is labelled with the different phases of insertion. Peak 1 is relative to the insertion of the loop and peak 2 is relative to the entry of the enlarged area. Mean (± s.d., *n* = 10) insertion forces are 32.5 ± 22.7 mN (peak 1) and 223.4 ± 92.2 mN (peak 2). **d**, Forces during extraction from the optic nerve. Several extraction trials (*n* = 10) are shown in grey while, in red, a selected example is labelled with the different phases of extraction. Peaks (3 and 4) are relative to the extraction of the flaps and the flat phase (5) is relative to the slippage of the loop through the nerve. Mean (± s.d., *n* = 10) extraction forces are 101.2 ± 36.2 mN (peak 3) and 100.3 ± 38.5 mN (peak 4). **e**, The different phases of insertion (1 and 2) are sketched. **f**, The different phases of extraction (3 to 5) are sketched.

### Visual stimulation

Visually-evoked cortical potentials (VEPs) have been characterized in New Zeeland White rabbits upon flash illumination (Fig. 4a). The peak amplitudes (PAs) and the peak latencies (PLs) of the major peaks present in the VEP (N1 and P1) have been measured using an electrocorticography (ECoG) electrode array (Fig. 4b). Each rabbit (*N* = 9) has been stimulated on both eyes (ipsilateral and contralateral with respect to the ECoG array) with light flashes of increasing luminance (0.1, 0.5, 1, 5, 10, and 30 cd s m^−2^). As expected the strongest and fastest response occurs for contralateral stimulation (Fig. 4c), since in rabbits 90 to 95 % of the fibers decussate at the level of the chiasma^36^. Similar to a previous report^37^, the average (± s.e.m.) P1 PL in contralateral stimulation (Fig. 4d) starts at 33.93 ± 2.59 ms for low luminance (0.1 cd s m^−2^) and reaches a plateau latency of 23.28 ± 1.86 ms for higher luminance (5 cd s m^−2^). Also, N1 PL in contralateral stimulation is comparable to previously reported data (29.41 ± 2.40 ms, 25.17 ± 2.17 ms, 24.46 ± 2.01 ms, 20.65 ± 2.79 ms, 19.80 ± 1.35 ms, and 20.00 ± 1.38 ms respectively for 0.1, 0.5, 1, 5, 10, and 30 cd s m^−2^).

**Figure 4.**
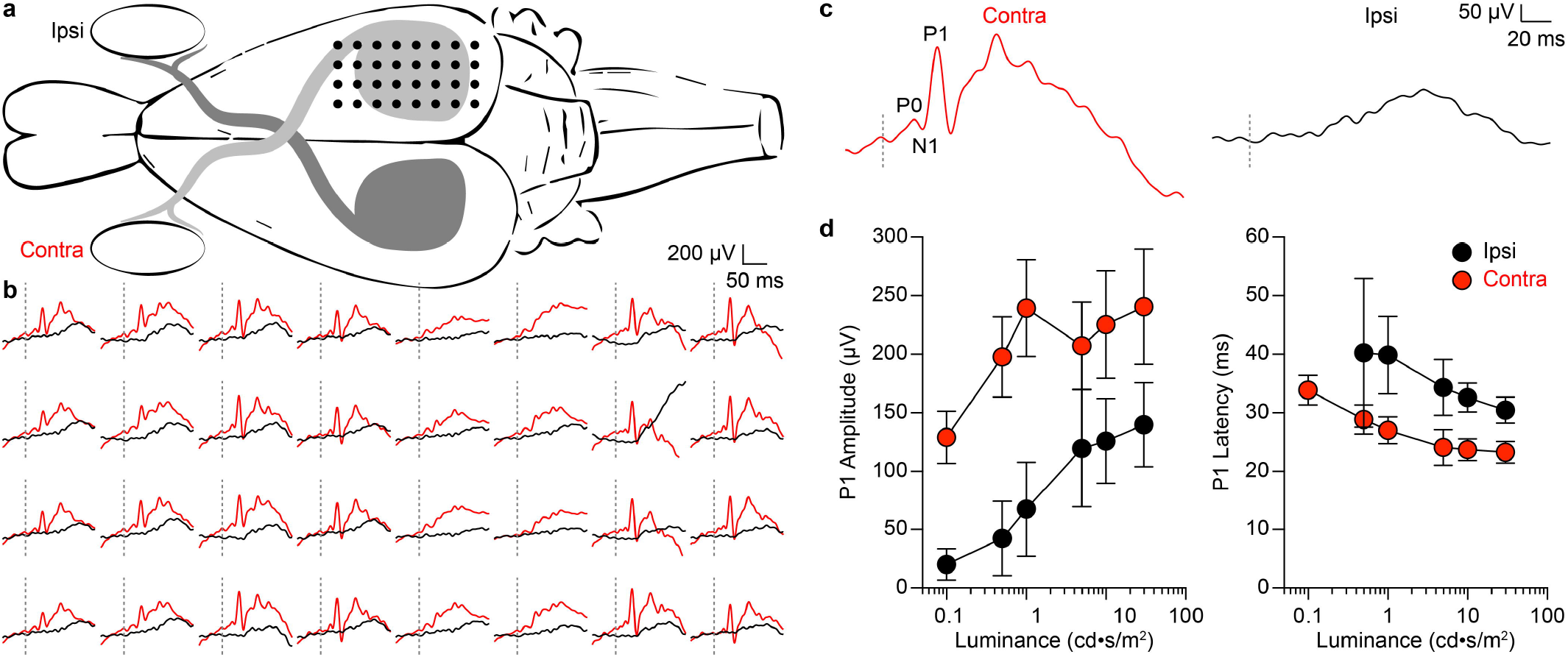
Visually-evoked cortical potentials. **a**, Recordings of VEPs have been performed with a ECoG array covering the visual cortex and a flash stimulation (4 ms, white LED) in the ipsilateral (black) or contralateral (red) eye. **b**, Example of traces (synchronous average of 10 responses) to a flash illumination of 0.5 cd s m^−2^ at the ipsilateral (black) or contralateral (red) eye. The dashed lines represent the occurrence of the flash. **c**, Example traces obtained from the average of the 32 recording channels (showed in **b**) for both ipsi- (black) and contra- (red) lateral stimulation. P0, N1 and P1 peaks are visible. The dashed lines represent the occurrence of the flash. **d**, Mean (± s.e.m.) of the P1 PAs (left) and PLs (right) with respect to the flash luminance (*N* = 9). P1 PAs and PLs have been measured in each rabbit from the average of the 32 recording channels.

### Electrical stimulation

The OpticSELINE has been implanted transversally in the optic nerve from the lateral to the medial side. Because of fiber decussation^33^, electrically-evoked cortical potentials (EEPs) have been measured with an ECoG array only in the contralateral visual cortex (Fig. 5a,b). Cathodic first asymmetrically balanced (1:5) electrical stimuli have been used. The ratio 1:5 has been previously demonstrated to be a good compromise between total pulse duration and stimulation efficiency^26^. In the same study, it has been also shown that placing the balancing anodic phase before the cathodic stimulation phase with a 1:5 ration has no influence. Moreover, we did not introduce any inter-phase gap since it has been demonstrated to have a significant effect only with symmetrically balanced stimuli^26^. To avoid fatigue in the nerve due to repetitive stimulations, only 6 electrodes of the OpticSELINE (1 to 6 in the top shank, from left to right) have been used in the first part of the study. In this condition, nerve fatigue due to repetitive stimulation has not been observed (**Supplementary Fig. 2**).

**Figure 5.**
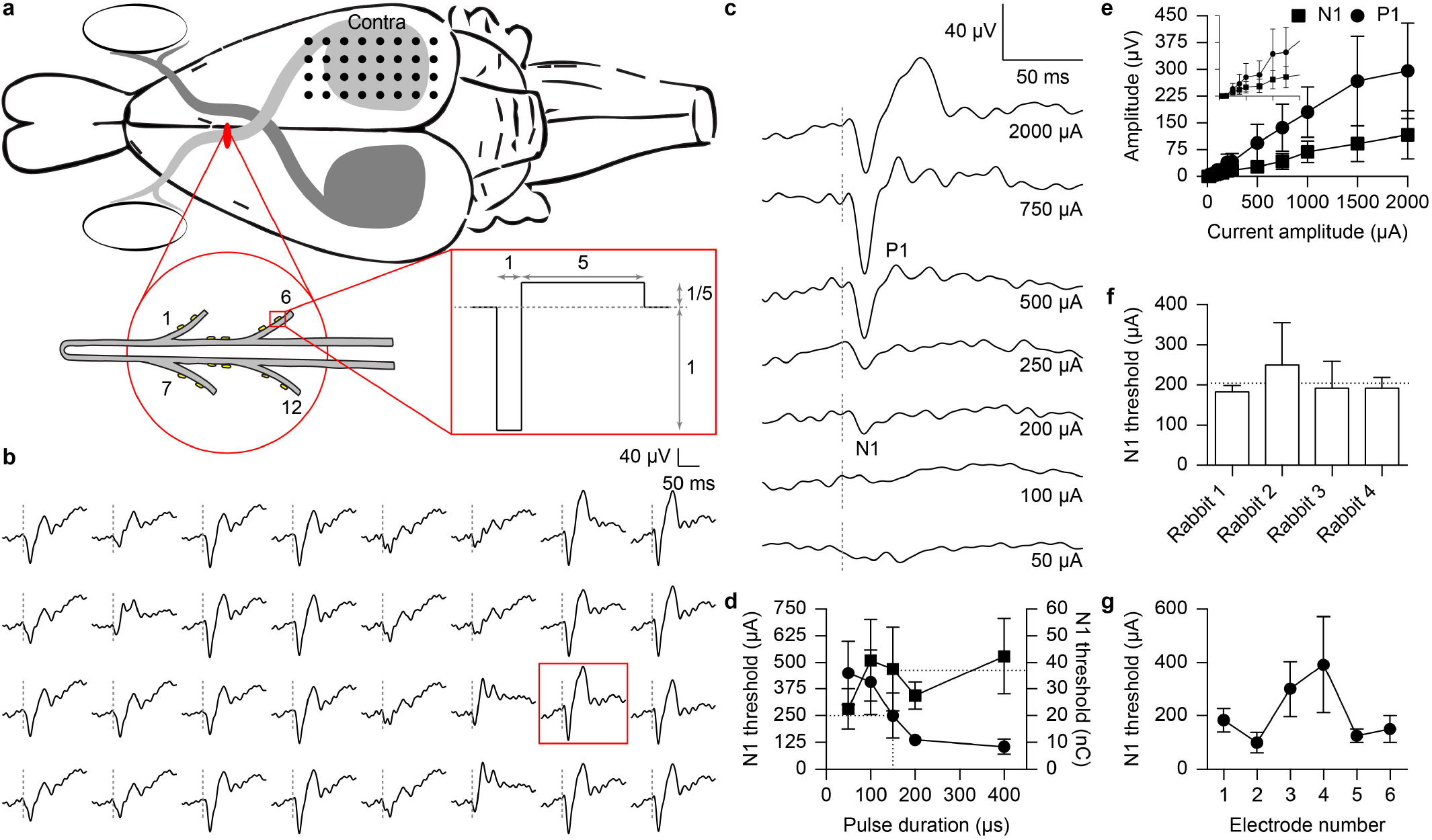
Electrically-evoked cortical potentials. **a**, EEPs have been evoked via the implanted OpticSELINE with biphasic current pulses. **b**, Example of traces (synchronous average of 10 responses) to a current pulse of 2 mA and 150 μs delivered from the electrode 6. The dashed lines represent the occurrence of the current pulse. **c**, Example of traces (synchronous average of 10 responses) to increasing current pulses of 150 μs from 1 recording electrode (red box in panel **b**). The appearance of N1 and P1 peaks is visible. **d**, Mean (± s.e.m.) N1 activation thresholds in *N* = 1 rabbit depending on pulse duration (circles, left y axis). The activation threshold is plotted also as charge delivered (squares, right y axis). The dotted lines show the activation threshold for current pulses of 150 μs, corresponding to 250 μA and 37.4 nC. **e**, Mean (± s.e.m.) amplitude of N1 (squares) and P1 (circles) for current pulses of 150 μs with respect to the stimulation amplitude (*N* = 4). The insert shows a magnification from 0 to 300 μA (x axis) and from 0 to 75 μV (y axis). **f**, Activation thresholds in *N* = 4 rabbits for current pulses of 150 μs (p = 0.86, Kruskal-Wallis). The dotted line represents the average threshold (204.17 μA, *N* = 4). **g**, Mean (± s.e.m.) activation thresholds (*N* = 4; p = 0.25, one-way ANOVA) depending on the electrode (from 1 to 6). In panels **d, e, f**, and **g**, the responses from the 32 recording electrodes have always been averaged. In panels **d, e**, and **f**, for each rabbit the quantifications from each of the 6 electrodes (from 1 to 6) have been averaged.

Under electrical stimulation of the optic nerve we found that both N1 and P1 peak amplitudes increase with the current amplitude of the stimulus (Fig. 5c,e). Then, we have measured the minimum current required to induce the N1 peak in the cortical response (average of the 32 electrodes of the ECoG). N1 current threshold decreases with the increase of the pulse duration; on the contrary, the pulse duration seems to have a minimal effect on the amount of charges required to activate N1 (Fig. 5d), as previously observed^38^. Therefore, the phase duration has been fixed to 150 μs. Under this condition, the mean (± s.e.m.) N1 threshold results in 204.17 μA (Fig. 5f). Interestingly, the electrodes on the main body (numbers 3 and 4) have a mean (± s.e.m.) threshold higher (Fig. 5g) than the electrodes on the flaps (numbers 1, 2, 5, and 6). This could be explained since in the pre-chiasmatic trait of the optic nerve the fibers with large diameter are localized more in the periphery of the nerve than in the center (**Supplementary Fig. 3**).

Retinal ganglion cells can generate action potentials up to few hundreds of Hz; for such reason, the artificial reproduction of this code requires high frequency electrical pulsing. We verified the possibility to use high frequency pulse trains (1, 2, 3, or 4 pulses at 1 kHz of repetition rate) for optic nerve stimulation (Fig. 6a). By increasing the number of pulses within the train (from 1 to 4) at constant current amplitude, the mean (± s.e.m.) N1 threshold is progressively reduced down to 46.15 ± 11.79 % (for 4 pulses) of the threshold obtained with a single pulse (Fig. 6b). Similarly, the cortical activation (e.g. P1 PA) can be modulated by increasing the number of pulses instead of changing the current intensity (Fig. 6c).

**Figure 6.**
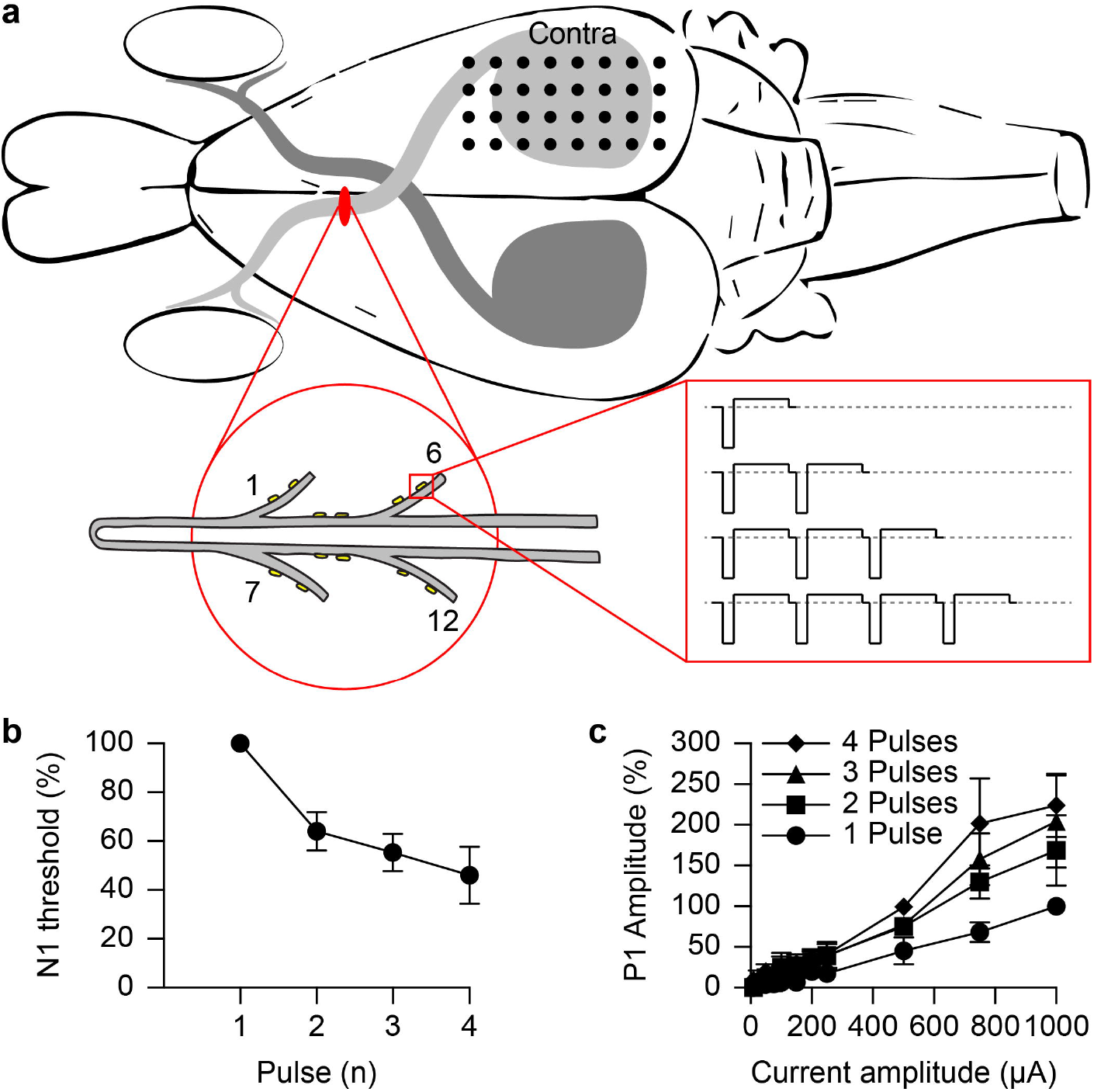
Electrically-evoked cortical potentials with pulse trains. **a**, EEPs have been evoked via the implanted OpticSELINE with biphasic current pulses arranged in packages containing 1, 2, 3, or 4 pulses at 1 kHz of repetition rate. **b**, Normalized mean (± s.e.m.) of N1 activation threshold with stimuli composed of 1 to 4 pulses (*N* = 4). The normalization has been performed with respect to the stimulus with 1 pulse. **c**, Normalized means (± s.e.m.) of P1 PA (*N* = 4) for trains with 1, 2, 3, or 4 pulses (circles, squares, triangles, and diamonds respectively). The normalization has been performed with respect to the amplitude for 1 pulse at 1000 μA. In panels **b**, and **c**, the responses from the 32 recording electrodes have always been averaged. Moreover, for each rabbit the quantifications from each of the 6 OpticSELINE electrodes (from 1 to 6) have been averaged.

### Stimulation selectivity

Last, we investigated the spatial selectivity of intraneural stimulation of the optic nerve with single pulses. A blind source separation approach has been used to quantitatively extract differences between the EEPs resulting from the stimulation through different electrodes of the OpticSELINE. Independent component analysis (ICA)^39^ has been chosen since it is steadily gaining popularity among blind source separation techniques to disentangle information linearly mixed into multiple recorded data channels so as to prepare multivariate data sets for more general data mining^40–42^.

We hypothesized that each EEP is composed of ‘shared’ components and ‘meaningful’ components: the former ones are characterized by similar time courses regardless of the stimulating electrode activated in the OpticSELINE, while the latter ones have time courses that are specific to one specific stimulating electrode (or to a small subset at maximum). ICA has been performed on the 32 ECoG recordings in order to highlight the presence of meaningful components that may have dispersed in the common cortical signal. ICA linearly projects the 32 original time courses into 32 new maximally-independent time courses, here called independent components (ICs), as weighted sums of the original time courses. The region of the visual cortex in which each IC is present has been determined by plotting the activation map (see Methods) of each IC on the 32 electrodes of the recording array (Fig. 7a). In addition, ICs have been classified in different categories, based on their time course (**Supplementary Fig. 4**), namely: artifact (i.e., containing the stimulation artifact), noise (i.e., containing a high frequency signal), flat (i.e., not containing any peaks in their waveform), common (i.e., having a similar time courses for each of the stimulating channels), and meaningful (i.e., having different time courses depending on the stimulating channel). In a representative example (*N* = 1 rabbit, current amplitude of 750 μA), amongst the 32 ICs, 26 have been labelled as meaningful, 2 as common, 1 as flat, 1 as noise, and 2 as artifact (Fig. 7b, **Supplementary Fig. 5**). In this representative example 9 stimulating channels out of 12 resulted in a meaningful cortical activation, while 3 of them induced flat responses only. By back-projecting only the meaningful ICs onto the original channel space, the original data can be also filtered to highlight only the meaningful component (**Supplementary Fig. 6**). By increasing the current amplitude of the pulse, the number of ICs classified as meaningful increases, it reaches his maximum value (26) at 500 μA until 1000 μA, and then decreases (**Supplementary Fig. 7b**). This is probably due to the beginning of the saturation of the cortical response; therefore, more ICs are classified as common instead of meaningful.

**Figure 7.**
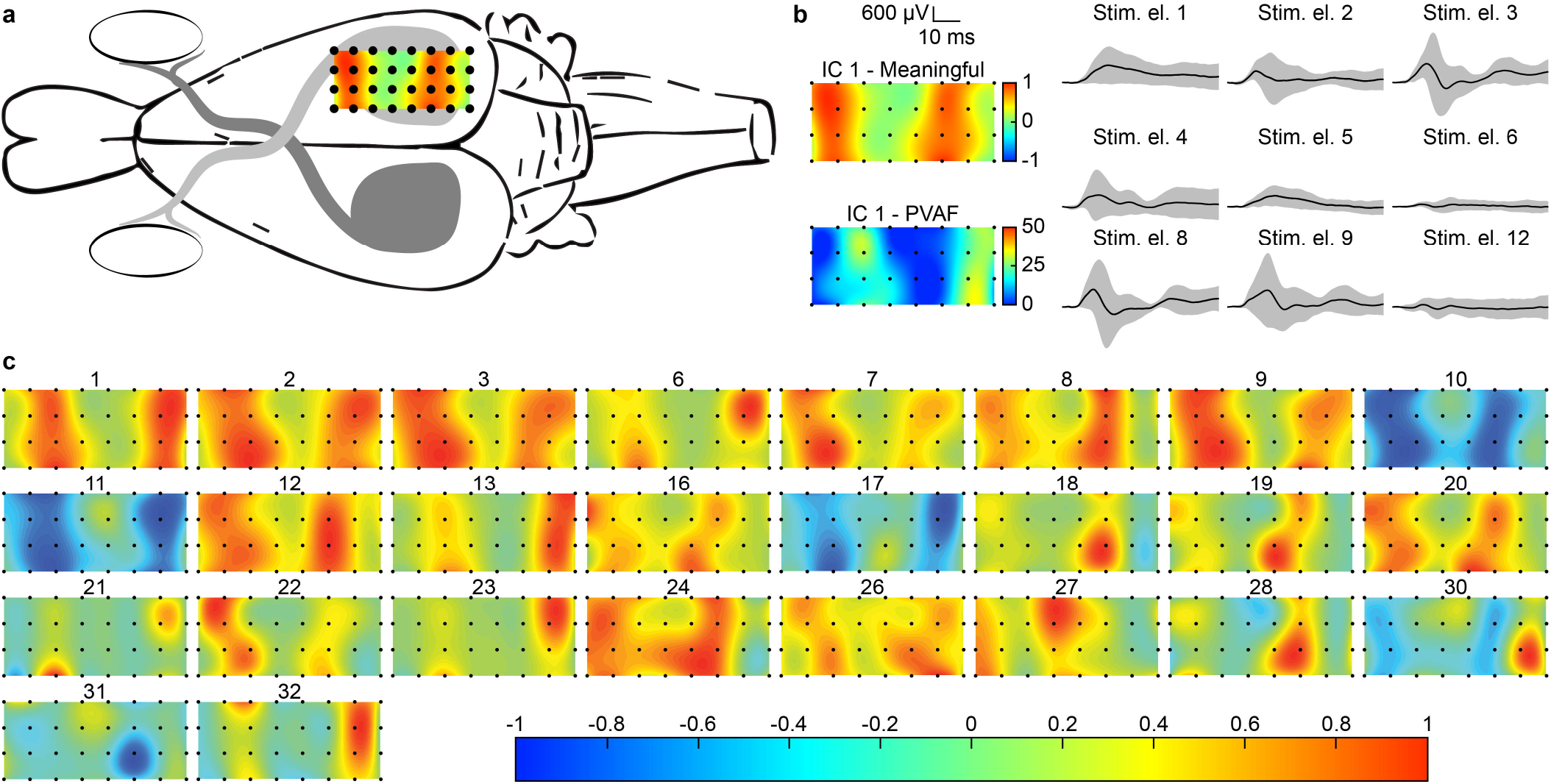
Activation maps of the meaningful ICs. **a**, Example of an activation map corresponding to IC 1 projected in correspondence of the ECoG array. **b**, Example of the activation map, the PVAF, and the time courses of a representative meaningful component (IC 1). A time course is shown for each stimulating electrode of the OpticSELINE. **c**, The 26 meaningful ICs are present in distinct regions of the visual cortex. Representative example from *N* = 1 rabbit with a current amplitude of 750 μA.

The activation maps of the 26 meaningful ICs show that they are present in distinct regions of the visual cortex, which suggests that the components of the original signal are spatially segregated in the visual cortex (Fig. 7c). In addition, each meaningful ICs exhibits a different degree of activation depending on the stimulating electrode used. To quantify this, for each meaningful IC, the peak-to-peak amplitude of the early portion of the time courses (from 5 to 25 ms after the pulse) induced by each stimulating electrode was computed and normalized amongst all the stimulating electrodes. All of the 9 stimulating electrodes have at least one meaningful IC whose activation is maximized when stimulating through this particular electrode. Furthermore, by interpolating the contribution of every stimulating electrode to each IC, we found their distribution map within the optic nerve (Fig. 8a and **Supplementary Fig. 8**). The area corresponding to 90 % of the activation level (red lines) is confined to a small area around this electrode, or in few cases it spreads over 1 or 2 neighboring electrodes (Fig. 8b). The spatial segregation of the ICs at the level of the visual cortex (Fig. 7) and the high localization of ICs in correspondence of the OpticSELINE electrodes (Fig. 8) are considered indirect signatures that the different stimulating electrodes recruit different population of optic nerve fibers, thus confirming that the OpticSELINE allows selective visual cortex activation.

**Figure 8.**
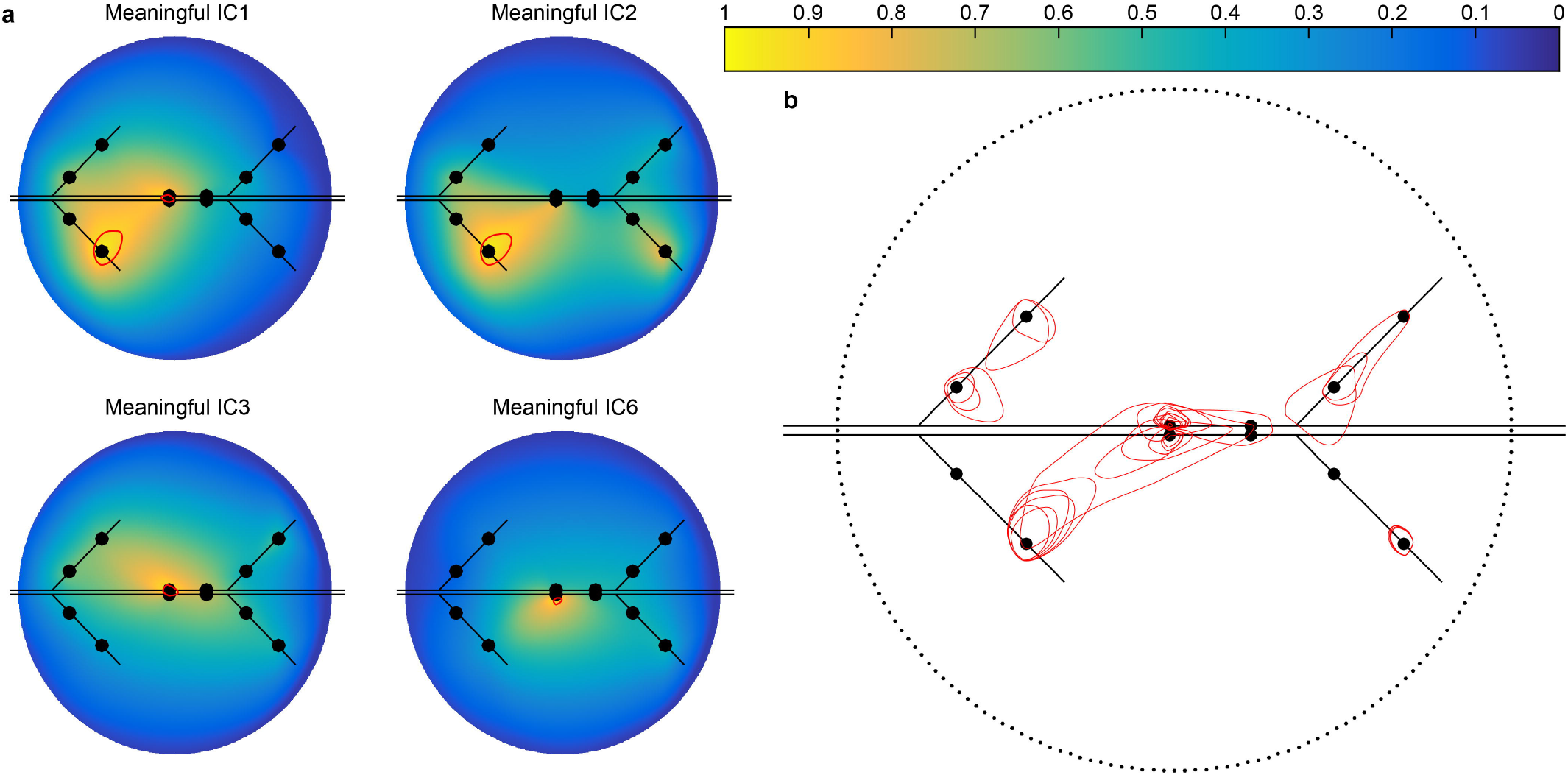
ICs distribution map at the level of the optic nerve. **a**, Example of the distribution maps of the first 4 meaningful ICs within the optic nerve. Enclosed in the red contour is the region of the optic nerve which activation level is larger than 90% of the maximal activation. **b**, Overlay of the 90% activation contours (red) of all the meaningful ICs. The dotted circle represents the optic nerve. The sketch of the OpticSELINE is in black in both **a** and **b**. Representative example from *N* = 1 rabbit with a current amplitude of 750 μA.

## Discussion

Currently, the research on visual prostheses is largely focused on the development of novel retinal prosthesis^15–17,43–47^, either subretinal, epiretinal, or suprachoroidal. Nevertheless, beside the early pioneering works, some research groups are now attempting the stimulation of downstream regions of the visual pathway^12^. Among those, optic nerve prostheses aim at stimulating the axonal fibers from retinal ganglion cells during their pathway towards the optic chiasma. Following two initial studies on two blind patients affected by Retinitis pigmentosa^19,20^, optic nerve stimulation is now under validation in a clinical trial by another group which is targeting the optic disk in the retina^48^.

In the first clinical trial, the two patients have been implanted with epineural cuff electrodes, which are less invasive than intraneural arrays. However, epineural electrodes mostly target external fibers, while intraneural electrodes can stimulate also the central area of the nerve and provide higher selectivity^31^. In addition, intraneural electrodes are mechanically more stable than epineural^32^, therefore allowing a more reproducible and stable stimulation. For this reason, we selected an intraneural approach, confirming mechanical stability, due also to the lateral flaps, and selectivity in fiber recruitment as demonstrated by the blind source separation analysis.

In our experimental design, we choose to target the intra-cranial trait of the optic nerve. Epineural electrodes have been previously implanted in both the intra-orbital^20^ and intra-cranial trait of the nerve^19^. The comparison of the two clinical studies showed that intra-orbital stimulation induces smaller EEPs with respect to intra-cranial stimulation, while latencies are not significantly different; in other words, the perceptual threshold is higher for intra-orbital stimulation^9^. In addition, the intraorbital trait of the optic nerve must accommodate the eye movements; therefore, an intraneural electrode may be subjected to high cyclic strain which may later lead to failure. This is minimized in the intra-cranial trait. A second drawback of the intra-orbital placement of an intraneural electrode is due to the presence of the central retinal vein and artery entering the nerve approximately 1 cm far from the eye bulb; therefore, an intraneural placement may risk to damage them. Therefore, intracranial stimulation by intraneural electrodes seems more appropriate than intra-orbital stimulation. Compared to retinal prostheses, and in particular epiretinal implants, optic nerve stimulation differs since it activates only axonal fibers. During epiretinal stimulation, electric pulses may induce a direct activation of retinal ganglion cell fibers or an indirect activation via the internal retinal circuit. It is known that brief (hundreds of μs) cathodic epiretinal stimulation preferentially excites the axons of retinal ganglion cells, while pulses longer than 1 ms excite both retinal ganglion cells and bipolar cells^16,49,50^. A recent study has demonstrated that the increase of the stimulation to 25 ms per phase significantly reduces the unwanted activation of axon of passage. In turns, this enhances the indirect activation or retinal ganglion cells via the internal retinal circuits, which provides a more focal activation avoiding streak responses^51^. The disadvantage of long stimulations (25 ms per phase) is the reduction of the frame rate available. Currently the Argus II operates in the range of 3 to 60 Hz^52^; however, the increase of the pulse duration to 25 ms may limit the maximum frame rate to about 10 Hz. An attractive strategy for visual prostheses is neuromorphic encoding^53–55^, where the conversion between an image and the pattern of electrical stimulation delivered to the tissue is based on an in-silico morphing of the retinal process. In this case the stimulation pattern delivered to the neurons (e.g. retinal ganglion cells) can be a representation of the natural code. Retinal ganglion cells can fire action potentials up to few hundreds of Hz, therefore the reproduction of this natural code may require a high temporal precision. It is now evident that an epiretinal strategy may not combine the use of long pulses (e.g. 25 ms per phase) with a neuromorphic approach. On the contrary we have demonstrated that optic nerve stimulation can be implemented at high frequency (i.e., 1 kHz) thus allowing the reproduction of the natural code of retinal ganglion cells with high temporal fidelity. Optic nerve stimulation appears as good strategy for neuromorphic prostheses or for the testing of neuromorphic algorithms in animal models.

An important aspect to enable the OpticSELINE as visual prosthesis is the capability to induce selective optic nerve stimulation and generate spatially organized phosphenes. The first clinical trial with epineural electrodes already demonstrated that optic nerve stimulation may induce spatially organized phosphenes^22^. The C-Sight project demonstrated in rabbits that the position of the cortical channel with the highest amplitude can be spatially modulated by applying current-steering methods to the intraneural stimulation of the intra-orbital trait of the optic nerve^25^. We have found that with an intraneural multi-electrode array is possible to associate ‘meaningful’ cortical activation patterns with specific electrodes of the array, confirming that the OpticSELINE induces activation of the visual cortex in a reproducible and spatially organized manner. This is possible because of the selective activation of nerve fibers, which could open the possibility to elicit behaviorally relevant visual activation patterns by optimizing the stimulation protocol. This represents a fundamental breakthrough towards the implantation of the optic nerve stimulation to restore functional vision.

## Acknowledgement

This work has been supported by École Polytechnique Fédérale de Lausanne, Medtronic, Bertarelli Foundation, and Wyss Center for Bio and Neuroengineering. Dr. Artoni is supported by the European Union’s Horizon 2020 research and innovation programme under Marie Skłodowska Curie grant agreement No. 750947 (BIREHAB).

## Author contributions

V.G. designed the stimulation protocol, performed data analysis and blind source separation. A.C. designed and fabricated the OpticSELINE and performed mechanical and electrochemical characterizations. P.V. performed in-vivo and histological experiments. F.A. performed data analysis and conceived the blind source separation approach. S.A.R.P. participated in the design of the stimulation protocol and in data analysis. D.L.D.P. participated in the design and microfabrication of the OpticSELINE and performed mechanical characterizations. S.M. designed the study and supervised the activities related to electrode development and the blind source separation approach. D.G. designed the study, led the project, and wrote the manuscript. All the authors read, edited, and accepted the manuscript.

## Methods

### Electrode microfabrication

The OpticSELINE was developed by using micro photolithography and thin-film techniques. A silicon wafer was used as sacrificial layer. After the cleaning of silicon wafer (10 min Acetone; DI rinse; 10 min Isopropanol), two layers of polyimide PI2610 (HD MicroSystems) were spun on the substrate (2000 rpm for 30 s). Samples were hard baked in an oven with nitrogen flux at 350 °C for 1 hr. LOL and S1813 (Microposit) were spin-coated on the wafer respectively at 1000 rpm for 20 s and 4500 rpm for 30 s. The substrate was exposed by using a glass photomask at a dose of 180 mJ cm^−2^. The sample was developed in MF319 for 30 s and rinsed in DI water. A layer of titanium (20 nm) and gold (250 nm) were sputtered on the substrate and a lift-off technique was used to release the pattern of traces, active sites and pads (overnight immersion in remover 1165). Two layers of PI2610 were spun on the substrate (2000 rpm for 30 s) and hard baked in the oven with nitrogen flux at 350 °C for 1 hr. An aluminum mask (200 nm) was deposited on the substrate by thermal evaporation. S1813 was spun on the sample and exposed (glass photomask, 180 mJ cm^−2^) and the wafer was developed in MF319 for 30 s. S1813 was removed (2 min in remover 1165) and dry etching was used to etch the excess of polyimide (40 sccm of O_2_; 150 W, 1 hr). The aluminum mask was etched away and electrodes were peeled off from the wafer. The three-dimensional geometry was conferred to the OpticSELINE by securing the device on a stainless-steel mold. The mold has 4 holes in correspondence of the 4 flaps; a needle was used to secure each flap inside the hole. Alignment holes were included on the mold to ease the placement of the device. A thermal treatment (1 hr, 200 °C) was used to memorize the curved shape of the flaps. The 3D devices were connected to the polyimide-based extension cable: a silver conductive glue was used to connect the corresponding pads of the two elements (Ablestik-Henkel; 1 hr at 130 °C). Then the flexible extension cable was connected to a printed circuit board (PCB) by silver conductive glue. A surgical needle with a looped wire (Ethicon) was inserted through the device. Flexible wires were soldered to the PCB and then a linear Omnetics connector was attached. Two-component biocompatible silicone (Silbione-Bluestar Silicones) was applied on all the soldering.

### Electrochemical characterization

Cyclic Voltammetry was performed using a three-electrode setup immersed in phosphate-buffered saline (PBS) solution and applying a 10-cycle potential ramp at a scan rate of 1.5 V s^−1^ between −0.5 V and 0.5 V. Impedance measurement was performed using a three-electrode setup immersed in PBS solution and applying a sinusoid of 10 mV between 100 Hz and 100□kHz. Accelerated aging test was performed for 6 days at temperature of 87 °C with an accelerating factor of 32. Different OpticSELINE arrays have been immersed in glass beakers filled with PBS solution; the beakers were sealed and stored in oven.

### Mechanical tests

Both experiments were performed using the same setup composed of a press to secure the nerve, a 10 N load cell, and an explanted rabbit optic nerve. During insertion experiments, first the nerve was pierced by the needle, then the electrode was pulled at a constant speed of 15 mm min^−1^ to insert the device inside the nerve. During extraction experiments, first the device was implanted inside the nerve, then the electrode was pulled at a constant speed of 15 mm min^−1^ to completely extract the device from the nerve. In both cases, insertion and extraction forces were measured by a load cell.

### Animal handling and surgery

Animal experiments were performed under the animal authorization GE1416. Female New Zealand White rabbits (> 16 weeks, > 2.5 kg) were sedated with an intramuscular injection of xylazine (5 mg kg^−1^). Anesthesia and Analgesia were provided with an intramuscular injection of an anesthetic mix composed by: medetomidine (0.5 mg kg^−1^), ketamine (25 mg kg^−1^), and buprenorphine (0.03 mg kg^−1^). If required, anesthesia was prolonged with a second injection (half dose) of the anesthetic mix. Eye drops were placed on the eye to prevent eye drying. The rabbit was placed on a heating pad at 35°C for the entire procedure. Oxygen was provided with a mask to prevent hypoxia during the anesthesia. The head was shaved and cleaned with 70% ethanol and betadine. The rabbit’s head was then secured gently within a stereotactic frame (David Kopf Instruments). Prior to cortical skin incision, a mix of lidocaine (6 mg kg^−1^), bupivacaine (2.5 mg kg^−1^), and epinephrine (0.1 mg kg^−1^) was injected subcutaneously on the surgical sites. After 5 minutes, the skin was opened and pulled aside to expose the skull; finally, the skull was cleaned with cotton swabs. First a temporal craniotomy was made to access the left optic nerve. The OpticSELINE was inserted in the left optic nerve from lateral to medial in the pre-chiasmatic area. Then a second craniotomy was made to expose the right visual cortex. A32-channel epi-dural ECoG array (E32-1000-30-200; NeuroNexus) was placed on the visual cortex. All rabbits were euthanized at the end of the acute recording procedures, while still under anesthesia, with an intravenous injection of pentobarbital (120 mg kg^−1^).

### Optic nerve anatomy

To determine the average nerve diameter, optic nerves (*n* = 10) were explanted from 5 female New Zealand white rabbits (> 16 weeks, > 2.5 kg), immediately embedded in the optimum cutting temperature compound (O.C.T. Tissue-Tek, Qiagen), and frozen at −20 °C. 10-μm sections were obtained with a cryostat (Leica Microsystems) and mounted on glass slides. Images were taken with a slide scanner (VS120-L100, Olympus).

For the myelin staining, the optic nerve was extracted after surgery and fixed overnight in PFA 4%. The tissue was then dehydrated with increasing concentrations of ethanol and embedded in paraffin. 5-μm thick sections were cut using a microtome (Leica Microsystems) in the portion close to the optic chiasma. The Woelcke staining for myelin was performed on the sections after dewaxing by specialized technicians and the images acquired in transmitted light at 40X magnification and analyzed using ImageJ.

For the myelin basic protein staining, antigen retrieval in citrate buffer (pH 6) was performed on dewaxed sections. The samples were then blocked for one hour in PBS-Triton 0.1 % + NGS 5 % (Jackson Immuno Research) and incubated at 4 °C overnight with the primary antibody (rabbit anti-MBP 1:200, ab40390, Abcam). The following day, the sections were washed in PBS and incubated 1 hour at room temperature with the secondary antibody (goat anti rabbit Alexa-488). The samples were mounted for imaging using Fluoromount (Sigma-Aldrich) solution. The images were acquired with a confocal microscope (LSM-880, Carl Zeiss) at 63 X magnification and analyzed using ImageJ.

### Electrophysiology

For optic nerve stimulation the OpticSELINE was attached to a current stimulator (IZ2MH; Tucker-Davis Technologies), while for cortical recordings the ECoG array (E32-1000-30-200; NeuroNexus) was connected to an amplifier (PZ5; Tucker-Davis Technologies) via a 32-channels analog headstage (ZIF-Clip^®^ Analog Headstage; Tucker-Davis Technologies). Optic nerve stimulation was performed with 13 pulse amplitudes (10, 25, 50, 75, 100, 150, 200, 250, 500, 750, 1000, 1500, and 200 μA) and 5 pulse durations (50, 100, 150, 200, and 400 ms) delivered in a scrambled manner. Data were filtered between 0.5 Hz and 2 kHz and digitalized at 12 kHz. Epochs (from −100 to 750 ms) synchronous to the onset of the stimulation were then extracted from the data stream and data analysis was performed with Matlab (Mathworks).

### Blind source separation

For each recording electrode and each stimulation intensity, epochs were concatenated and processed with an AMICA^48^ core and GPU-processed infomax reliable ICA (RELICA) algorithm^49^. Dimensionality reduction on the data as a preprocessing step to ICA was not performed^50^. RELICA allowed to test the repeatability of ICs appearing in decompositions of bootstrapped versions of the input data and to retain only stable ICs for further analysis. Given the multivariate dataset from the 32 recording electrodes Xfelectrode, time), ICA extracts an unmixing matrix W (32 x 32) such that the IC time courses S=WX are maximally independent. Rows of W represent the weights applied to each electrode to obtain the corresponding ICs *S*. The column i^th^ of the mixing matrix A (pseudoinverse of W) represents the weight of the i^th^ IC on each recording electrode and can be represented as an activation map. Each activation map was obtained by projecting the weights of the unmixing matrix A onto the layout of the ECoG array, then by spatially interpolating them with a spline-function, and finally by normalizing the maps to the maximal absolute value present in the interpolated map. IC grand average time courses, obtained by performing the average over trials for each stimulation intensity, formed the IC-EEPs. The percent of variance accounted for (PVAF) by each IC on each electrode was computed and represented as a PVAF activation map. ICs were categorized into several classes, namely: flat, common, artifact, noise, and meaningful (**Supplementary Fig. 4a**). First, low frequency components of the signal were removed using a zero-phase high-pass filter with a 5 Hz cut-off frequency. Then artifact ICs, exhibiting a large activation within the first 5 ms from the stimulus onset, were identified by visual inspection and removed. To verify that the artifact ICs were correctly identified, all the other ICs were back-projected. The initial portion of the back-projected signal (0 to 5 ms) was indeed exclusively affected by the manual removal of the artifact ICs when compared to the original signal (**Supplementary Fig. 9**). Following the identification of the artifact, the noise ICs were identified by computing the frequency plot of the signal; the ICs exhibiting unusual peaks in the 250 to 500 Hz frequency range were labeled as noise (**Supplementary Fig. 4b**). Amongst the remaining ICs (**Supplementary Fig. 4c**), the ones with a peak-to-peak variation (Δ) in the time course (time frame from 5 to 25 ms after the stimulus) smaller than 3 times the standard deviation of the time course (σ) were labelled as flat (Δ/σ < 3). To separate meaningful ICs from common ICs, a similarity index was used; that is the correlation between the time courses of the different stimulating electrodes. ICs were classified as common (i.e. with visible meaningful activation but with similar time courses for all the stimulating electrodes) for a mean similarity index larger than 0.4. Distribution maps in the optic nerve were obtained by interpolating with a spline function the contribution of every stimulating electrodes to each IC. The contour of the optic nerve was set to zero.

### Statistical analysis and graphical representation

Statistical analysis and graphical representation were performed with Prism (GraphPad Software Inc.). The normality test (D’Agostino & Pearson omnibus normality test) was performed in each dataset to justify the use of a parametric or non-parametric test. In each figure p-values were represented as: * p < 0.05, ** p < 0.01, *** p < 0.001, and **** p < 0.0001. Data are reported as mean ± s.e.m. or mean ± s.d., *n* is used to identify the number of electrodes used; *N* is used to identify the number of animals.

### Data availability

The authors declare that all other relevant data supporting the findings of the study are available in this article and in its Supplementary Information file. Access to our raw data can be obtained from the corresponding authors upon reasonable request.

